# Development of modular geminivirus-based vectors for high cargo expression and gene targeting in plants

**DOI:** 10.1101/2024.06.29.601216

**Authors:** Matthew Neubauer, Katie Vollen, Jose T. Ascencio-Ibanez, Linda Hanley-Bowdoin, Anna N. Stepanova, Jose M. Alonso

## Abstract

Viral vectors can be useful tools for expressing recombinant proteins as well as delivering gene-editing machinery. Despite their utility, the development and subsequent optimization of these tools is often a difficult and tedious process. Thus, although considerable work has been done to create useful viral vectors for gene editing and protein expression, there is a lack of understanding of how best to design these vectors for specific applications. For instance, it is often unclear whether the inclusion of heterologous promoter sequences or different viral components will improve cargo expression or replicon accumulation. To address some of these hurdles, we designed a GoldenBraid (GB)-compatible viral vector system based on the geminivirus – Beet curly top virus (BCTV). This system allows for simple, modular cloning of a variety of reporter constructs. Making use of this modular cloning strategy, we compared a variety of alternative viral vector architectures. Interestingly, native BCTV promoters outperformed the constitutive *35S* promoter, while the removal of the BCTV virion-sense genes promoted reporter expression. Intriguingly, these modifications had no effect on total replicon accumulation. These results show the utility of the new modular BCTV-based viral vectors for protein expression and gene targeting applications, as well as uncover design principles that may inform future geminivirus-based viral vector architectures. We anticipate that the availability of this new modular system will spark the broad adoption of replicon-based strategies in protein expression and gene editing experiments in plants.

## Introduction

Viral vectors have proven useful in various biotechnology applications, such as genome editing and protein expression. Geminivirus-based viral vectors have been used for the delivery of genome editing enzymes, such as zinc finger nucleases (ZFNs) and Cas9, as well as repair templates (RT) for homology-directed repair (HDR) (Butler et al., 2016; Wang et al., 2017; Yu et al., 2020; Gil-Humanes et al., 2017; Dahan-Meir et al., 2018, Eini et al., 2022). Geminiviruses are a family of pathogenic viruses with circular, single-stranded DNA genomes that collectively have a wide host range and agricultural impact (Zerbini et al., 2017). The ability of geminivirus-based viral vectors to accumulate at high levels in plant cells has been shown to be critical for providing the high levels of repair template (RT) necessary for successful Gene Targeting (GT) in plants (Baltes et al., 2014).

Beet curly top virus (BCTV), a member of the geminivirus family, is a curtovirus with a monopartite genome that consists of virion-sense and complementary-sense open reading frames (ORFs) and an intergenic region (IR) (Figure 1A). Like other geminiviruses, BCTV replicates by rolling-circle replication (Hanley-Bowdoin et al., 2013). The complementary-sense ORFs encode the Rep (C1), C2, Ren (C3), and C4 proteins. The complementary-sense ORFs are separated from the virion-sense ORFs by the short AT-rich IR that contains the origin of replication and viral promoters for bidirectional transcription. The virion-sense ORFs encode the coat protein (CP, V1), a movement protein (MP, V2), and V3. V2 acts as a suppressor of host silencing and is required for systemic infection (Luna et al., 2020). The virion-sense ORFs are located directly downstream of the IR, which contains conserved late promoter elements (CLEs) that can be activated following infection (Hur et al., 2008). Of all the DNA elements in the geminivirus genome, only Rep and *IR* are absolutely required for replication (Gutierrez et al., 1999; Hanley-Bowdoin et al., 2000), while Ren affects replication efficiency (Settlage et al., 2005). For most biotechnological applications, efficient replication of the geminivirus-derived vectors is all that is desired. In typical applications, the ability to move between cells and support different aspects of the virus life cycle conferred by the other viral genes is typically not required or desirable in these types of vectors. Although the genome size of fully functional viruses is constrained by movement requirements (Gilbertson et al., 2003), these constraints are likely removed when the cell-to-cell movement and encapsidation of the replicon are no longer of interest. However, the size limits of a stable and highly replicative geminivirus-derived vector are not known.

**Figure 1.**
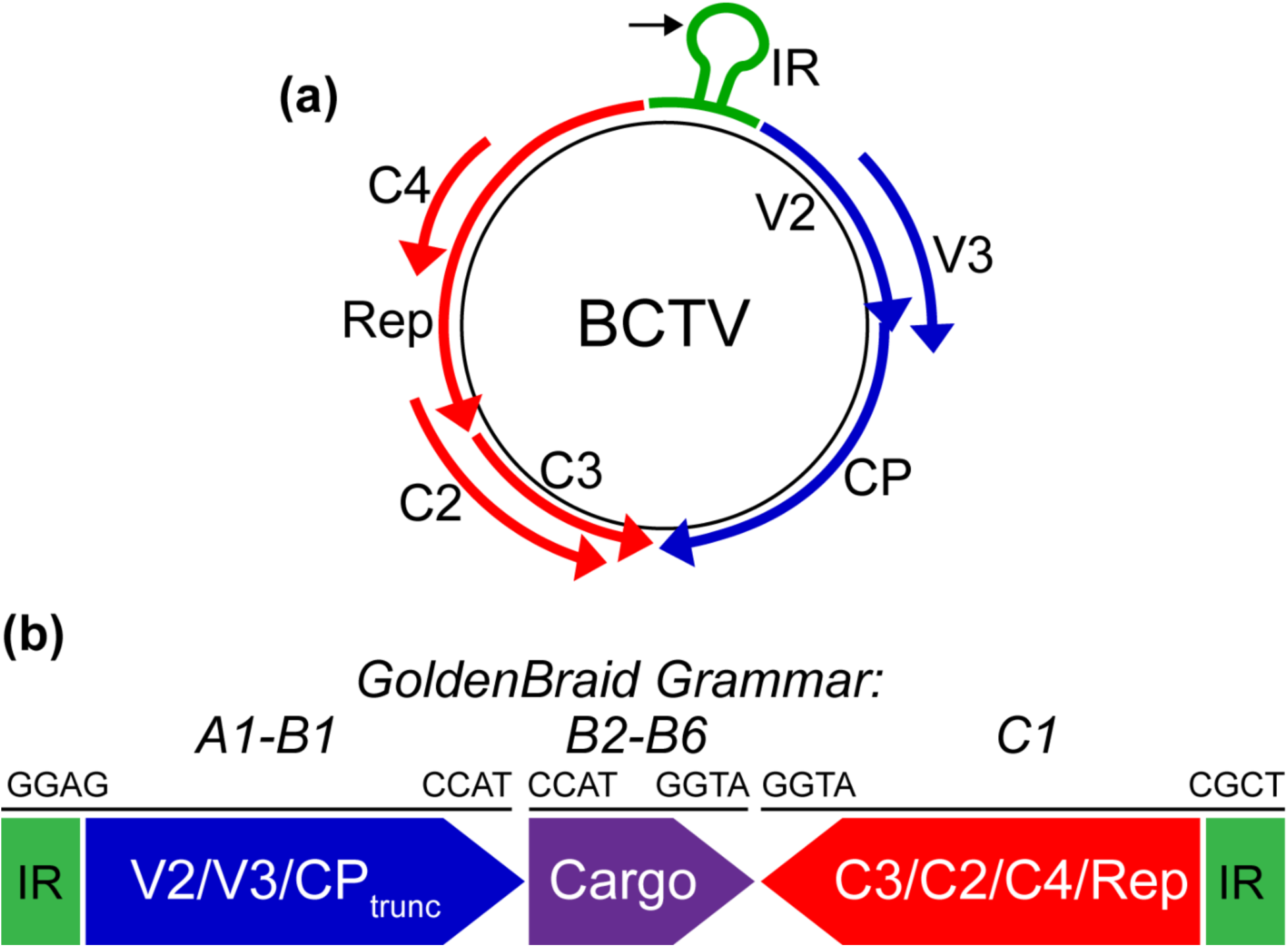
**A.** Diagram of the BCTV genome, which is composed of virion-sense (blue) and complementary-sense (red) genes and an intergenic region (IR, green). The black arrow denotes the location of the nick site near the nonanucleotide sequence that is required for rolling circle replication. **B.** GoldenBraid cloning was implemented to generate genetic elements containing portions of the BCTV genome. The virion-sense (blue) and complementary-sense (red) genes were cloned with A1-B1 and C1 grammar, respectively, while the cargo (purple) contained B2-B6 grammar. The 4-nt overhangs at the 5’ and 3’ ends of each genetic element are illustrated.

Various design approaches have been used to generate BCTV-based viral vectors, including removing the virion-sense genes and inserting heterologous promoters (Kim et al., 2007; Golenberg et al., 2009; Chung et al., 2011; Eini et al., 2022). For instance, a BCTV-based viral vector was utilized for Cas12a-mediated gene-editing in plants (Eini et al., 2022). To facilitate the expression of *Cas12a* and other cargoes, the authors removed the virion-sense genes and inserted a Cauliflower mosaic virus (CaMV) *35S* promoter upstream of the cargo (Eini et al., 2022). Various heterologous promoters were tested for compatibility with BCTV-based vectors including CaMV *35S*, the Cassava vein mosaic virus promoter, and the Arabidopsis *UBIQUITIN3* and *ACTIN8* promoters, with a *GFP* reporter employed to monitor cargo expression levels (Kim et al., 2007). In another study, the authors sought to induce gene silencing and inserted the gene of interest (GOI) in the antisense orientation downstream of a truncated form of the virion-sense ORFs (Golenberg et al., 2009).

The BCTV *IR* contains three 30-bp conserved late element (*CLE*) motifs, which act to drive expression of the virion-sense genes. These elements alone were shown to be capable of driving the expression of a *GUS* reporter gene in transgenic Arabidopsis lines (Hur et al., 2008). Expression of the reporter was particularly strong in actively dividing tissues and in seedlings.

The transcriptional activity of the *CLE* elements was found to diminish as the plant aged. However, infection of mature plants with BCTV restored reporter gene expression. This demonstrates that although the *CLE* elements can mediate high levels of gene expression, this activity is also greatly impacted by the presence of other BCTV components either directly or indirectly through the changes induced on the host cell cycle (Hur et al., 2008).

GoldenBraid (GB) cloning is based on the use of Type IIS restriction enzymes and modular DNA parts that contain standardized ’grammar’ of 4-nt overhangs (Sarrion-Perdigones et al., 2011; Sarrion-Perdigones et al., 2013). A major benefit of GB cloning is that it enables existing genetic elements, such as promoters, terminators, and tags, to be assembled in the desired order in a single tube reaction and with great efficiency. This allows for previously generated parts to be reused in new constructs without the need to subclone them from scratch. Furthermore, the standardized grammar allows for parts to be easily shared between labs.

In this study, we sought to determine the optimal composition of a BCTV-based viral vector for the delivery of repair template in gene targeting experiments and to enable high levels of expression of a GOI. Utilizing the GB cloning approach, we generated and tested a variety of viral vector designs. The inclusion of a heterologous promoter upstream of a GOI was found to be unnecessary, with the intrinsic *IR* sequence acting as a stronger promoter than the constitutive CaMV *35S*. The BCTV viral vector was able to accommodate cargo that is 4 kb in size without a reduction in replicon accumulation. Successful GT events were detected following the delivery of gene editing machinery by the BCTV-based viral vector. Removal of the BCTV virion-sense genes greatly increased cargo protein expression but did not enhance GT efficiency. These observations demonstrate the utility of tailoring viral vector design depending on its intended use. Furthermore, the new GB-based BCTV-derived tools developed here should facilitate the flexible design and testing of more efficient and application-tailored viral vectors.

## Materials and Methods

### Strains, vectors, and seed lines

GB cloning vectors, including the pUPD2 entry vector and pDGB3alpha and pDGB3omega vectors, were obtained from the Orzaez lab (Sarrion-Perdigones et al., 2011, 2013). The BCTV viral sequences used in this study were amplified from a BCTV-Logan infectious clone (provided by the Hanley-Bowdoin lab, originally from the D. Bisaro Lab), and cloned into the pUPD2 entry vector. The *Nicotiana tabacum GUS-NPTII* transgenic line (Wright et al., 2005) and pLSLZDR plasmid were gifts from Dr. Dan Voytas. Wild-type *N. benthaminana* was grown from the lab stock that originated from seeds provided by Dr. Devarshi Selote.

### Plant Growth Conditions

*N. tabacum* and *N. benthamiana* seeds were surface-sterilized in a solution containing 10% bleach and 0.05% Tween for 1 hour, washed three times with sterile dH_2_O and plated on 0.5X MS plates. One-week-old seedlings were transferred to soil (Sungro Metromix 830) in 24 cell flats and grown at 23°C with 60% humidity under a 16-h-light and 8-h-dark cycle. Four-week-old plants were used in transient expression experiments.

### Plasmid Construction

GB cloning was employed to generate clones used in this study (Sarrion-Perdigones et al., 2011). BCTV sequences were subcloned using a BCTV-Logan infectious clone as a template. The full-length virion-sense portion of the genome was amplified using the primers BCTV IR A1F and BCTV 5’ +Stop B2R (Table 1). To construct virion-sense deletion (*ΔVS*) clones, the left border-flanking *IR* region was amplified using the IRTrunc A1F and IRTrunc B1 R primers (Table 1). Amplification of the complementary-sense portion of the genome was performed using the BCTV 3’ C1F and BCTV 3’ C1R primers (Table 1). Amplification of the complementary-sense portion of the genome lacking the IR region was performed using the BCTV 3’ C1F and BCTV C4 C1R primers (Table 1). PCR products were purified and assembled individually into the pUPD2 entry vector.

**Table 1.**
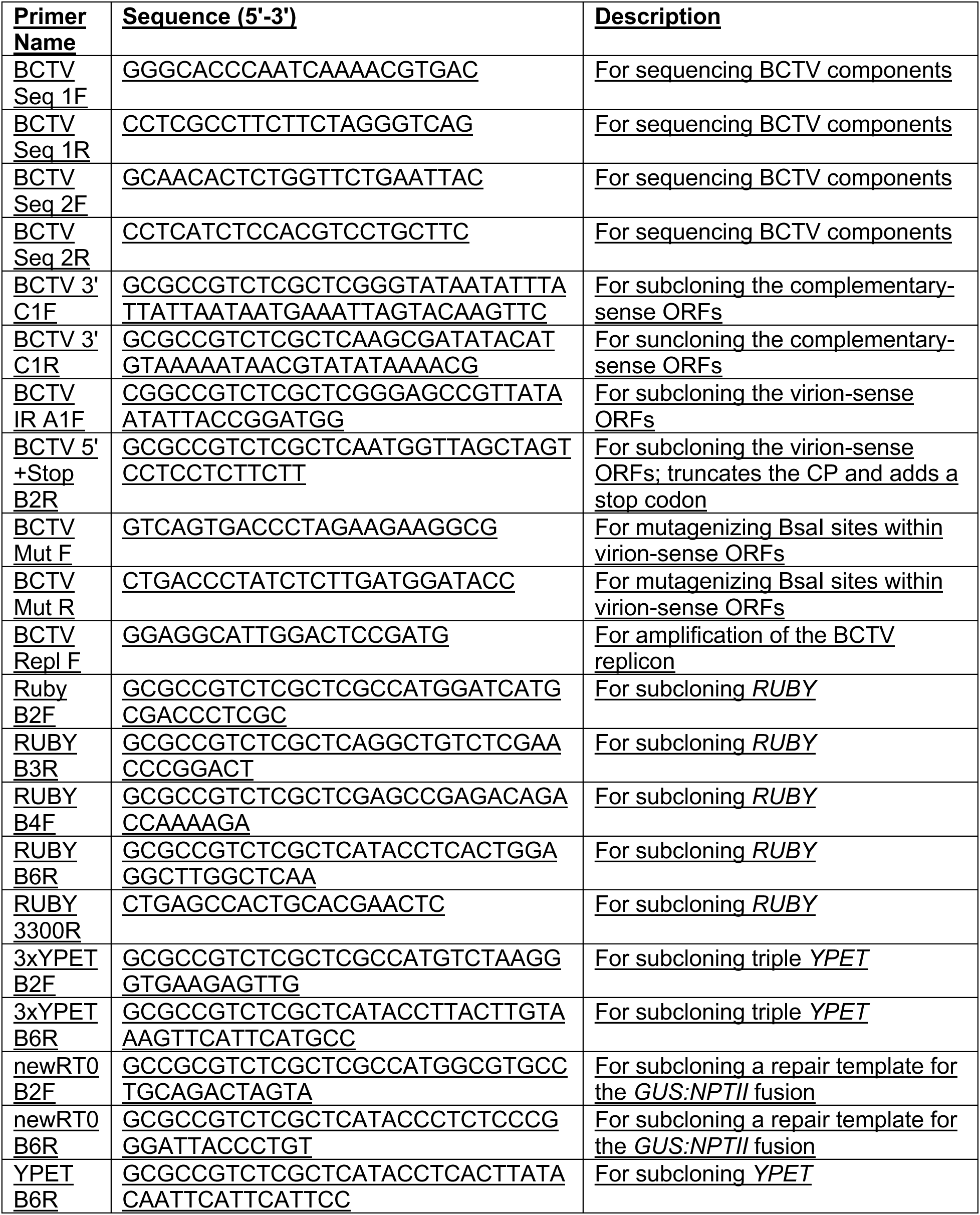
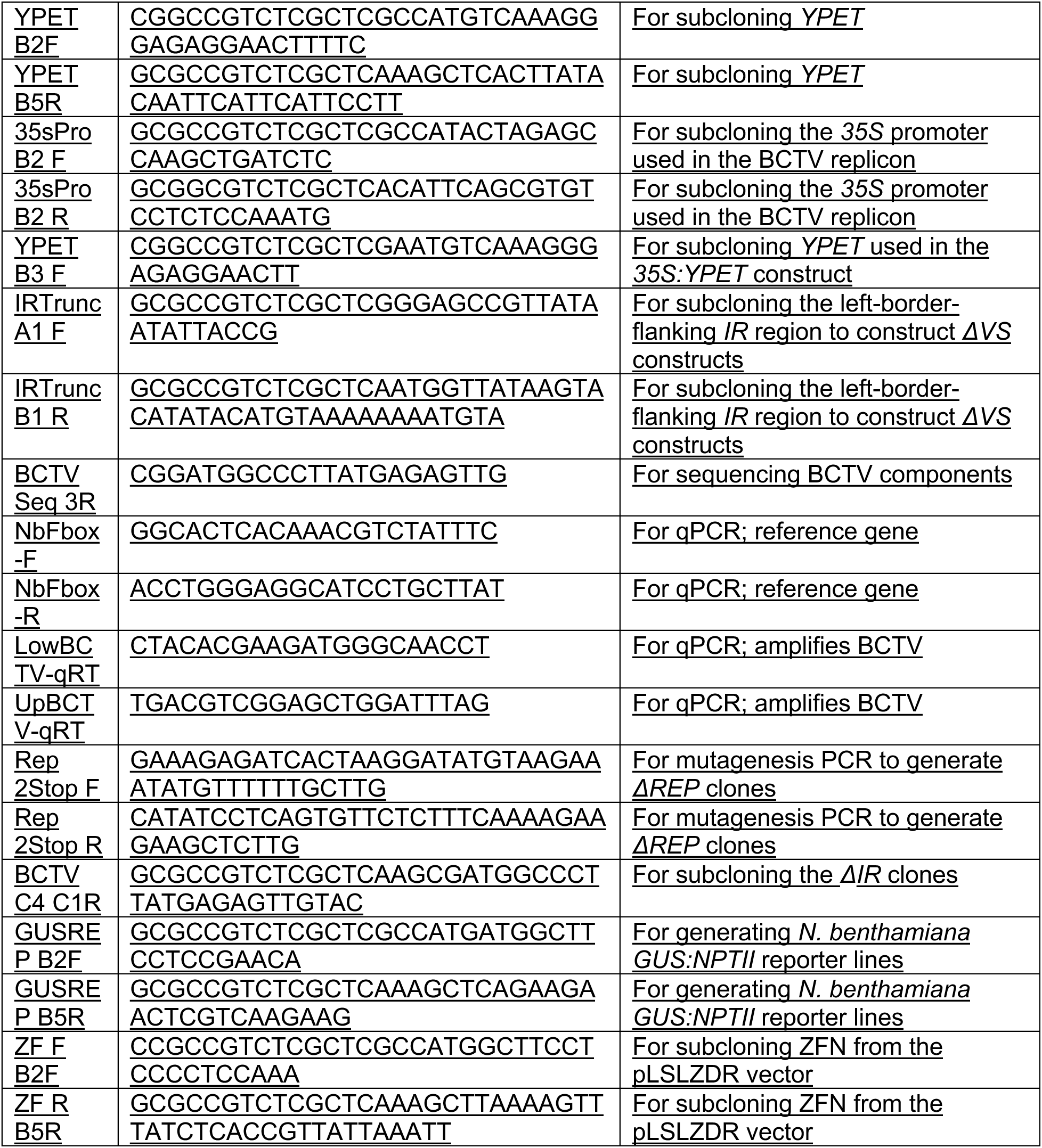
Primer Table.

For later assembly as BCTV cargo, *YPET*, *3xYPET*, *35S* promoter, gene targeting repair template, and *RUBY* sequences were domesticated with compatible 5’ and 3’ grammar sequences. *YPET* was amplified from an existing pDGB3alpha clone using the YPET B2F and YPET B6R primers, or, for placement downstream of the *35S* promoter, YPET B3F and YPET B6R primers (Table 1). *3xYPET* was amplified from an existing pDGB3alpha clone using the 3xYPET B2F and 3xYPET B6R primers (Table 1). The *35S* promoter sequence was amplified using the 35sPro B2F and 35sPro B2R primers (Table 1). For use as a gene targeting repair template, the repair template portion of the pLSLZDR plasmid was amplified using the newRT0 B2F and newRT0 B6R primers (Baltes, et al., 2014; Table 1). *RUBY* was cloned as two fragments using the RUBY B2F and RUBY B3R primer pair and the RUBY B4F and RUBY B6R primer pair (Table 1). The template used to amplify the *RUBY* reporter was the *35S:RUBY* plasmid from the Yunde Zhao Lab (He et al., 2020). The ZFN from the pLSLZDR plasmid was amplified using the ZF F B2F and ZF R B5R primers (Baltes et al., 2014; Table 1). Following amplification, these amplicons were assembled into the pUPD2 vector.

PCR-based site-directed mutagenesis was performed on pUPD2 entry clones to generate the BCTV virion-sense ORF and *ΔREP* complementary-sense clones (Qi and Scholthof, 2008). The BCTV Mut F and BCTV Mut R primers were used for the mutagenesis of a BsaI site within the BCTV virion-sense ORF. To generate *ΔREP* clones, stop codons were introduced into a pUPD2 clone containing the complementary-sense ORF with the REP 2STOP F and REP 2STOP R primers (Table 1). Sequencing of BCTV pUPD2 clones was performed using the BCTV Seq 1F, BCTV Seq 1R, BCTV Seq 2F, BCTV Seq 2R, and BCTV Seq 3R primers (Table 1). These pUPD2 clones were used in subsequent assemblies to generate pDGB3alpha clones.

Assembly reactions to generate pUPD2 entry clones and pDGB3omega clones were performed using the BsmBI-v2 TypeIIS restriction enzyme and T4 Ligase, New England Biolabs (NEB). Assembly of pDGB3alpha clones was performed using BsaI-HF and T4 Ligase (NEB).

Assembly reactions were performed using a thermocycler set to cycle between digestion and ligation reactions (40 cycles of 37°C for BsaI-HF or 42C for BsmBI-v2 for 5 minutes, followed by 16°C for 5 minutes).

Assembly of the pDGB3alpha clones *BCTV:YPET*, *BCTVΔREP:YPET*, *BCTVΔIR:YPET*, *BCTVΔVS:YPET*, and *BCTVΔVSΔREP:YPET* constructs was performed by assembling viral components previously assembled into pUPD2 as A1-B1 and C1 parts with a pUPD2 clone containing *YPET* cloned as a B2-B6 part. Assembly of *BCTV:35S:YPET* and *BCTVΔVS:35S:YPET* was performed using pUPD2 clones containing the *35S* promoter and *YPET* cloned as B2 and B3-B6 parts, respectively. The *35S:YPET* pDGB3alpha clone was created by assembling pUPD2 clones containing the *35S* promoter cloned as an A1-B1 part, *YPET* cloned as a B2-B5 part amplified using the YPET B2F and YPET B5R primers (Table 1), and the *35S* terminator cloned as a B6-C1 part.

Assembly of the *BCTV:3xYPET* pDGB3alpha clone was performed by assembling pUPD2 clones containing the BCTV viral components cloned as A1-B1 and C1 parts and the *3xYPET* reporter cloned as a B2-B6 part. Similarly, the *BCTV:RUBY* and *BCTV:ΔVS:RUBY* pDBG3alpha clones were generated using pUPD2 clones containing the *RUBY* reporter cloned as B2-B3 and B4-B6 parts.

To generate clones used for gene targeting experiments, the pDGB3alpha clone *35S:ZF* was assembled using pUPD2 clones containing the *35S* promoter, the ZFN, and the *35S* terminator cloned as A1-B1, B2-B5, and B6-C1 parts, respectively. The pDGB3alpha clones *BCTV:RT*, BCTV*ΔREP:RT, BCTVΔVS:RT,* and *BCTVΔVSΔREP:RT* were assembled using pUPD2 clones containing the viral components cloned as A1-B1 and C1 parts with the repair template cloned as a B2-B6 part. The pDGB3omega clones *ZFN + BCTV:RT* and *ZFN + BCTVΔVS:RT* were generated by assembling the corresponding alpha clones together.

#### Agroinfiltrations and Transient Expression

Plasmids were transformed into *Agrobacterium tumefaciens* strain GV3101 (pMP90) by electroporation with selection on Luria-Bertani (LB) plates containing 50 μg mL^−1^ kanamycin sulfate, 20 μg mL^−1^ gentamicin, and 10 μg mL^−1^ rifampicin. For agroinfiltration experiments, 5 mL liquid LB cultures were grown for 20 hours at 30°C. *Agrobacterium* cells were resuspended in agroinfiltration buffer (10mM MES, 150 μM acetosyringone, 10 mM MgCl2, pH 5.7) at OD_600_ 0.2. After 3 hours, *Agrobacterium* was infiltrated into leaves using a needleless 1 mL syringe. For experiments involving co-infiltration, strains were resuspended in agroinfiltration buffer at OD_600_ 0.2 and then mixed prior to infiltration. Infiltrated plants were kept in the dark for 16 hours following infiltration before being returned to normal conditions.

### Transgenic GUS-NPTII N. benthamiana

The construct used to generate transgenic *GUS-NPTII N. benthamiana* plants was built using pre-existing *N. tabacum GUS-NPTII* plants (Wright et al., 2005). The broken *GUS-NPTII* fusion reporter was amplified from genomic DNA using the GUSREP B2F and GUSREP B5R primers and cloned into pUPD2 with GB B2-B5 grammar (Table 1). This B2-B5 part was combined with the *35S* promoter and *35S* terminator in the pDGB3alpha1 vector. The binary vector used for *N. benthamiana* transformation was a pDGB3omega2 clone created by combining a plant kanamycin resistance selectable marker with the *35S:GUS-NPTII* reporter using BsmBI-v2.

Transgenic plants were generated using *Agrobacterium* transformation and subsequent kanamycin selection (Horsch et al., 1989). Ten transgenic lines were generated, and PCR analysis confirmed that they all contained the T-DNA insertion. Agroinfiltration with the *ZFN+BCTV:RT* construct was performed in the T1 generation to identify the lines that performed the best in GT experiments. All lines tested were found to be functional. However, lines 1 and 13 generally gave the best results and were used in subsequent experiments.

Transgenic *N. tabacum* and *N. benthamiana* seedlings harboring the *GUS-NPTII* reporter were selected on 0.5X MS plates supplemented with kanamycin at a final concentration of 50 µg mL^−1^.

### GUS Staining

GUS staining experiments were performed at 7 days-post-infiltration (dpi) using a modified protocol (Baltes et al., 2014). Agroinfiltrated leaves were detached and vacuum-infiltrated for 5 minutes with X-Gluc solution (10 mM phosphate buffer, 10 mM EDTA, 1 mM ferricyanide, 1 mM ferrocyanide, 0.1% Triton X-100, and 1 mM X-Gluc (GoldBio)). Vacuum-infiltrated leaves were placed in petri dishes containing X-Gluc solution and incubated in the dark at room temperature with shaking at 80rpm for the first 2 hours. Infiltrated leaves were then incubated at 37°C for 20 hours in the dark. After staining, X-Gluc solution was removed, and the leaves were rinsed with distilled water. Chlorophyll extraction was performed on stained leaf tissue by soaking leaves in 80% ethanol for up to one week. The ethanol solution was replaced at least once per day.

Stained leaves were imaged using an Olympus digital camera. Successful GT events were quantified by counting the number of blue spots per leaf.

### DNA Extraction

To extract DNA, leaf samples were ground in CTAB buffer (1.4 M NaCl, 20 mM EDTA, pH8, 100 mM Tris-HCl, pH8, 3% CTAB (cetyltrimethylammonium bromide, Calbiochem)). Tissue grinding was performed by shaking tubes containing leaf samples and 100 µL of 1-mm glass beads in a Vivadent dental shaker for 5 seconds. DNA was isolated from infiltrated leaves through chloroform extraction and DNA precipitation, followed by washing with 70% ethanol as described (Doyle and Doyle, 1987).

### Quantitative PCR

Measurement of replicon accumulation was performed using qPCR on total DNA. The endogenous *NbFbox* gene was used as a reference gene and was amplified using the primers NbFbox-F and NbFbox-R (Eini et al., 2022; Table 1). To measure BCTV replicon accumulation, the LowBCTV-qRT and UpBCTV-qRT primers were used (Luna et al., 2017; Table 1). For each experiment, three biological replicates were isolated from unique leaves, and three technical replicates were performed on each biological replicate. Reactions were performed using a SYBR Green Master Mix (Applied Biosystems) with a Step One Plus Real-Time PCR

System (Applied Biosystems). qPCR analysis was performed using the Pfaffl method to adjust for discrepancies in primer efficiency (Pfaffl, 2001).

### Fluorescence Microscopy

Whole-leaf fluorescence microscopy was performed using a Leica M205. Images were merged using Leica software. Fluorescence microscopy on smaller leaf sections was performed using a Zeiss AxioImager M.2 upright microscope. ImageJ was employed to quantify fluorescence intensity. For quantification of fluorescence intensity, three equal-sized regions within the area of infiltration of each construct were quantified. To correct for autofluorescence, three ‘blank’ measurements were taken from un-infiltrated leaf regions, and the average of these measurements was calculated and subtracted from each of the three fluorescence measurements for each construct. The average of these three corrected fluorescence measurements was then calculated.

### DNA Gel Blotting

DNA gel blotting was performed on DNA extracted from *N. benthamiana* leaves that were infiltrated with *Agrobacterium*. DNA was digested with SalI restriction enzyme (NEB) overnight. Gel electrophoresis was performed on digested DNA using a 1% agarose gel. A dry transfer to a positively charged nylon membrane was performed overnight. A PCR DIG probe synthesis kit (Roche) was used for visualization of the dsDNA probe. The DNA probe was generated by amplifying the target sequence from *N. benthamiana* tissue infiltrated with *Agrobacterium* carrying the *BCTV:YPET* construct using primers BCTV MutR and UpBCTV-qRT (Table 1).

## Results

### Designing a modular BCTV-based viral vector

We sought to develop a modular BCTV-based vector that would enable rapid testing of different cargos, as well as making targeted alterations to the BCTV machinery encoded within the viral vector. To do this, we utilized a GB-based approach in which the BCTV virion-sense genes were cloned as an A1-B1 part, the cargo was cloned as a B2-B6 part, and the complementary-sense genes were cloned as a C1 part to generate the GB BCTV vector (Figure 1B). This modular approach simplifies the implementation of different BCTV-based vector designs. For instance, many alternative cargoes can be efficiently tested while utilizing the same virion-sense and complementary-sense fragments. Additionally, modifications can easily be introduced to the virus genetic machinery to optimize the system. Since the GB cloning strategy makes use of both BsaI and BsmBI Type IIS restriction enzymes, the DNA cargos and BCTV sequences that are used must lack these restriction sites. This necessitated the elimination of a single BsaI restriction site present in the virion-sense ORF. A 2-nt substitution was introduced using site-directed mutagenesis to modify this sequence, resulting in a Leucine to Valine substitution at V3 amino acid 42, and leaving V2 unaffected.

### Testing new modular BCTV-based vectors

To test whether the modifications made to convert the original BCTV viral sequences into a GB modular vector affect the basic replicative functions of the virus, we generated a BCTV viral vector containing a *YPET* reporter gene, *BCTV:YPET.* This construct contains the full-length BCTV viral genome, but to prevent the virus particles from forming, a stop codon was inserted into the *CP* gene after the 30th amino acid codon (Figure 2A). The *YPET* reporter was inserted, without any promoter or terminator, immediately downstream of the *CP* gene. We anticipated that *YPET* expression would be driven by the BCTV promoter normally controlling the transcription of virion-sense CDSs (*V2/V3/CP*) as part of the polycistron. When tested side-by-side with a traditional *35S* promoter-driven *YPET* T-DNA construct (Figure 2A) in transient assays in *N. benthamiana*, the *BCTV:YPET* construct consistently gave higher levels of fluorescence (Figure 2B, Figure S2A). We repeatedly observed that the fluorescence in leaves expressing *BCTV:YPET* typically peaked roughly three to four days post-inoculation (dpi) compared with a peak at two days generated by a *35S:YPET* control construct delivered as a T-DNA (data not shown). This delay in expression suggested that replicon accumulation is required for the high levels of expression observed in the *BCTV:YPET* construct.

**Figure 2.**
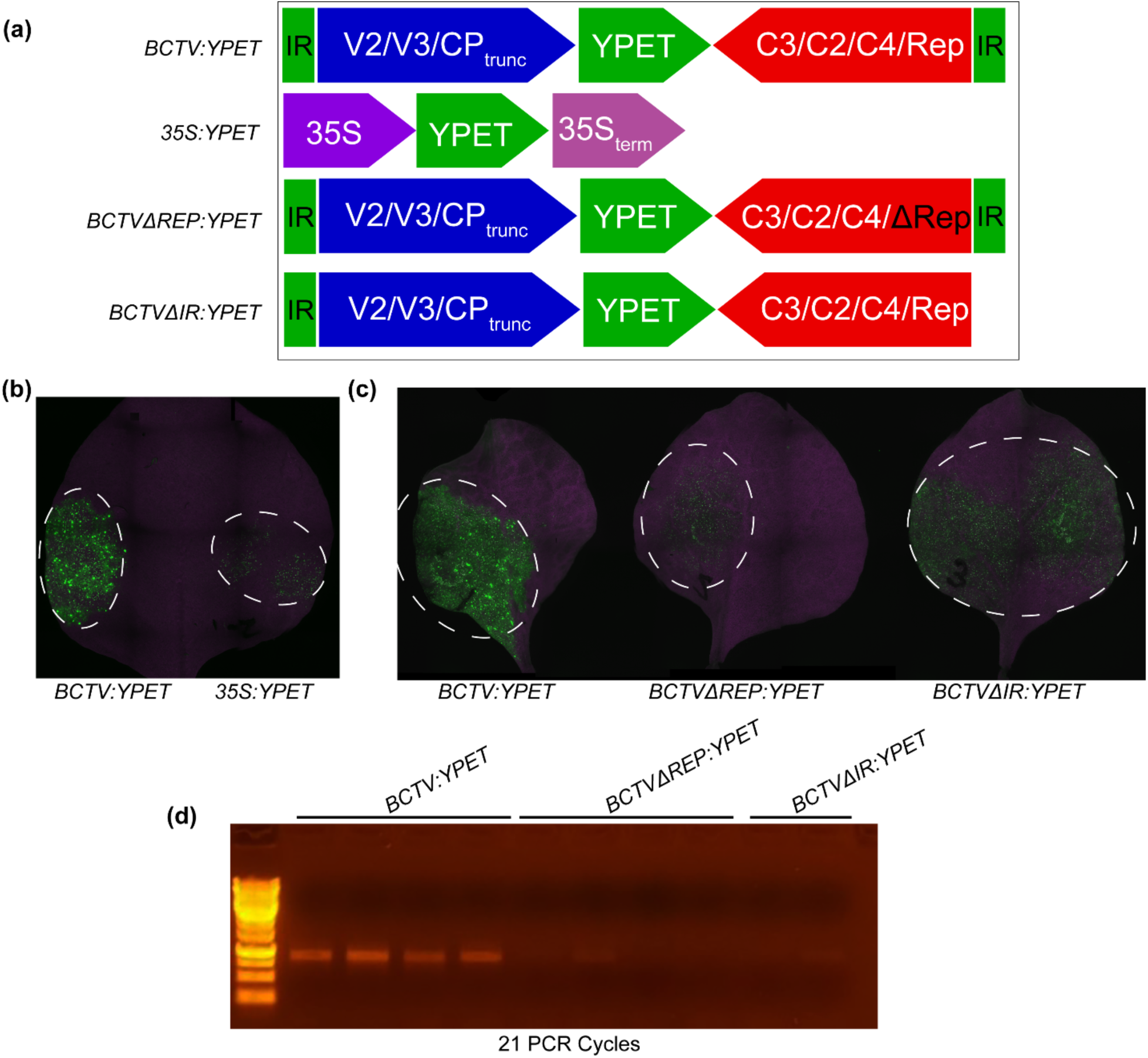
The BCTV viral vector mediates high levels of reporter expression which is dependent on the accumulation of the replicon. **A.** Diagram of the constructs used. A *35S*-driven *YPET* reporter was used as a control. **B.** YPET reporter expression from a BCTV-based viral vector was stronger than that resulting from a traditional *35S* promoter. Images were taken 5 dpi. **C.** Loss of *Rep* and the deletion of the right border-flanking downstream IR reduce reporter expression. **D.** PCR demonstrates that loss of *Rep* and the right-flanking IR motif inhibit replicon accumulation. PCR was performed using primers that can only amplify the circularized replicon. Each lane represents a unique biological replicate. DNA was extracted from plants at 5 dpi.

To demonstrate that the enhanced fluorescence observed following *BCTV:YPET* inoculation was due to viral replicon accumulation, and not just T-DNA-based expression, we generated two replication-deficient clones: *BCTVΔREP:YPET* and *BCTVΔIR:YPET* (Figure 2A). *BCTVΔREP:YPET* is identical to *BCTV:YPET*, except for two stop codons that were inserted into the 5’ end of *Rep*, mutating amino acids C20 and Q21 to stop codons (CAGTGT → TAGTGA). As Rep is known to be essential for initiation of viral replication, its inactivation is expected to abolish rolling circle replication. *BCTVΔIR:YPET* lacks the right border-flanking downstream *IR* which should also greatly inhibit the formation of the replicon. If the fluorescence observed with *BCTV:YPET* is replicon copy number-dependent, then these clones should exhibit significantly reduced fluorescence. Indeed, fluorescence mediated by the two replication-deficient clones was severely reduced relative to the *BCTV:YPET* clone (Figure 2C, Figure S2B). To verify that these sequence modifications indeed inhibit the formation of the BCTV replicon, PCR was performed on *N. benthamiana* tissue agroinfiltrated with these clones (Figure 2D). The PCR primers used to detect the replicon can produce a product using only the circularized form of the replicon and not the *Agrobacterium*-delivered binary vector (Figure S1). The lack of efficient PCR amplification suggests that the removal of *Rep* or the right border-flanking downstream *IR* significantly inhibits replicon accumulation and that this reduction likely contributes to the reduction in observed fluorescence (Figure 2C). Taken together, these results demonstrate that the modular, GB-compatible cloning approach is successful in producing functional BCTV-based viral vectors capable of carrying and expressing a promoterless cargo sequence and replicating *in planta*.

### Testing alternative vector designs

Despite the advantages of viral vectors as tools for plant biotechnology, little is known about how to best design these types of expression cassettes to maximize protein production. For example, in many plant viral expression vectors, the CaMV *35S* or other strong heterologous promoters are often used to drive the expression of the CDS of interest, despite the fact that many viral vectors may include native promoter elements known to drive high production of viral proteins. For instance, the BCTV *IR* is known to contain elements that mediate strong expression of virion-sense genes (Hur et al., 2007; Hur et al., 2008). Although our *BCTV:YPET* design incorporates the BCTV virion-sense genes, some previously published BCTV-based viral vectors lack these sequences (Eini et al., 2022). Although these sequences are not required for replicon formation and rolling circle replication, the BCTV virion-sense genes are involved in many processes, including movement of the virus and the regulation of the ratio of single-stranded (ss) to double-stranded (ds) viral DNA (Hormuzdi and Bisaro, 1993). Furthermore, the V2 protein inhibits host post-transcriptional gene silencing (Hanley-Bowdoin, et al., 2013; Luna et al., 2022). Therefore, it is possible that including these elements in the construct design will improve replicon formation or cargo expression.

To test the possible utility of native viral promoters in our modified GB BCTV vectors, we tested various construct architectures. In the first design, *BCTVΔVS:YPET*, a promoterless *YPET* CDS was placed between the left border-flanking *IR* and the *C3/C2/C4/Rep* sequences as an A1-B6 GB part, replacing the *V2/V3/CP_trunc_*. A version of this construct lacking a functional *Rep* was also included, *BCTVΔVS:ΔREP:YPET*. In these constructs *YPET* expression would be driven by the BCTV promoter normally controlling the transcription of virion-sense CDSs (*V2/V3/CP*) (Figure 3A).

**Figure 3.**
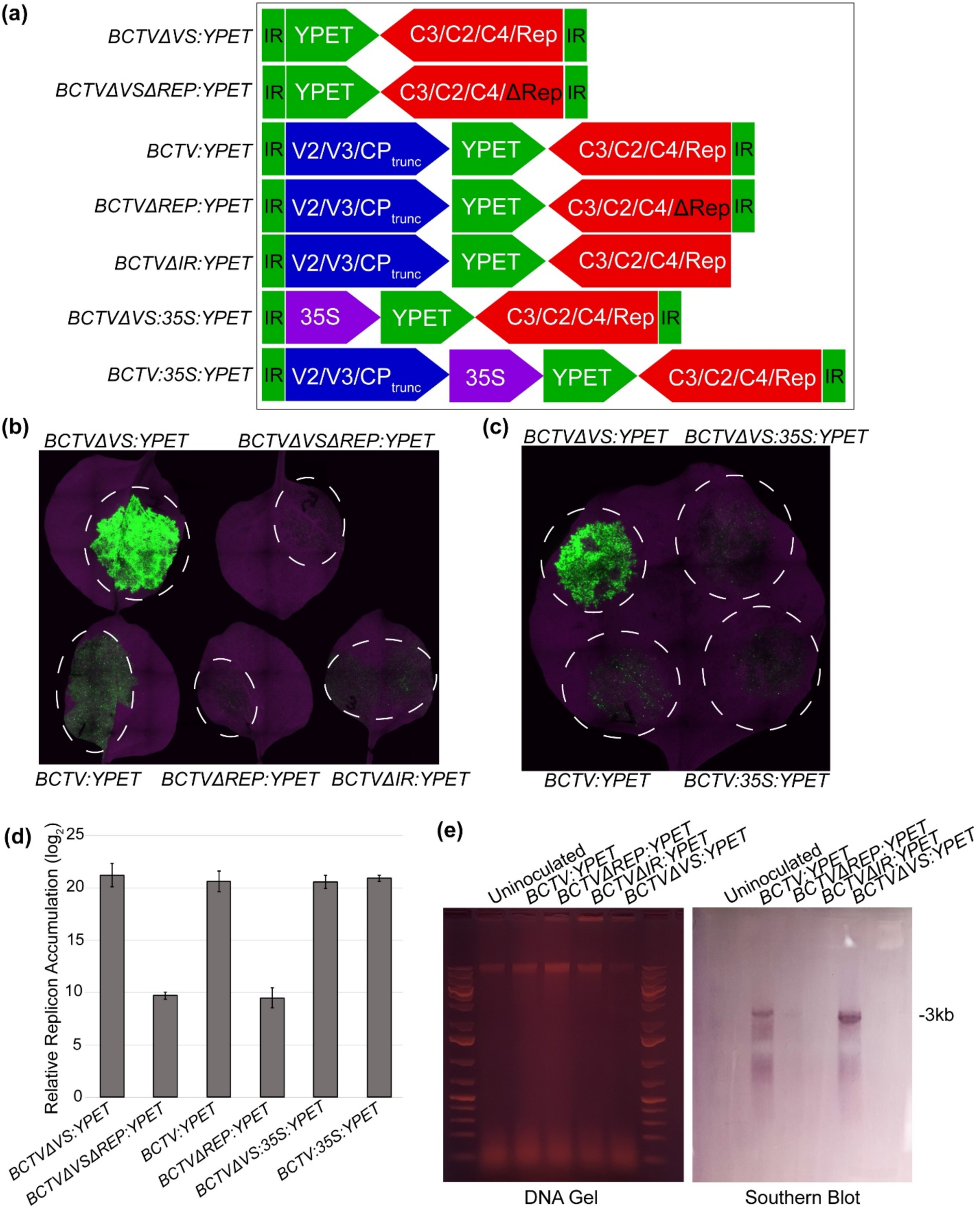
Removal of the BCTV virion-sense genes improves reporter expression, while inclusion of a heterologous promoter reduces expression. **A.** Diagram of various constructs used, including architectures that lack the virion-sense genes, *Rep*, and/or the right-flanking IR. **B.** Transient expression experiments were performed in *N. benthamiana* to determine how various viral vector architectures impact cargo expression. Images were taken at 5 dpi. The strongest expression was observed in leaves infiltrated with the *BCTVΔVS:YPET* construct, followed by the *BCTV:YPET* construct. **C.** Inclusion of the *35S* promoter upstream of the reporter reduces expression. YPET fluorescence resulting from constructs containing *35S* promoters was compared with those of constructs without heterologous promoters. BCTV viral vectors containing the *35S* promoter exhibited reduced fluorescence compared to those without. **D.** Insertion of the *35S* promoter upstream of the *YPET* reporter does not result in reduced replicon accumulation. qPCR was performed on DNA extracted from *N. benthamiana* leaves expressing various BCTV viral vector constructs. Amplification of DNA from constructs lacking *Rep* was significantly reduced, whereas constructs containing the *35S* promoter exhibited amplification levels similar to those without *35S.* **E.** A Southern blot was performed to detect BCTV replicon accumulation in infiltrated *N. benthamiana* leaves. The linearized ∼3kb dsDNA replicon was detected in leaves infiltrated with functional BCTV constructs. A lower molecular weight smearing pattern, consistent with the ssDNA form of the replicon, was also detected.

To test how removing the virion-sense CDSs would impact reporter expression, *Agrobacterium*-mediated transient expression experiments were performed in *N. benthamiana.* Leaves were imaged 5 days post-inoculation. In *N. benthamiana* transient assays, we observed that the *BCTVΔVS:YPET* construct consistently demonstrated higher reporter expression than the *BCTV:YPET* construct retaining the virion-sense genes (Figure 3B, Figure S2C). The loss of *Rep* in the *BCTVΔVSΔREP:YPET* construct severely reduced the YPET fluorescence, as was observed previously with the *BCTVΔREP:YPET* and *BCTVΔIR:YPET* vectors (Figure 3A), indicating that the high expression levels require replicon accumulation (Figure 3B, Figure S2C).

Additional architectures were tested to determine whether heterologous promoters would impact cargo expression. In one such design, *BCTV:35S:YPET*, a *35S:YPET* transcriptional unit was used as the GB B2-B6 part placed downstream of virion-sense genes (*V2/V3/CP_trunc_*) (Figure 3A). In parallel, a similar architecture was constructed, *BCTVΔVS:35S:YPET,* in which a *35S:YPET* transcriptional unit was placed directly downstream of the left border-flanking *IR* in place of the virion-sense genes (Figure 3A). To test whether insertion of a heterologous *35S* promoter could enhance reporter expression, these constructs were evaluated using *Agrobacterium*-mediated transient assays (Figure 3C). We found that the constructs containing the *35S* promoter performed worse than their counterparts without the *35S* promoter (Figure 3C, Figure S2D). This reduction in reporter expression was particularly evident in constructs lacking the virion-sense genes.

These results demonstrated that removal of the virion-sense genes has a positive impact on reporter expression, while the inclusion of the *35S* promoter upstream of the reporter has a detrimental impact. These changes in reporter expression could be the result of altered replicon accumulation. To test this, qPCR assays were performed on DNA extracted from *N. benthamiana* leaves infiltrated with *Agrobacterium* strains carrying various BCTV-based viral vector constructs (Figure 3D). qPCR demonstrated that despite mediating significantly different levels of reporter expression, the *BCTV:YPET*, *BCTVΔVS:YPET*, *BCTV:35S:YPET*, and *BCTVΔVS:35S:YPET* constructs supported similar levels of replicon accumulation. These results indicate that the differences observed in reporter gene expression between these four constructs are not the result of changes to replicon accumulation and could instead be due to some other factors, such as ratios of ss and ds replicon or different transcriptional activity of the promoter architectures tested.

As mentioned above, one explanation for the differences in reporter expression mediated by the various BCTV viral vectors is differences in the ratio of ds and ss replicon that accumulates within the cell. Although BCTV replicons exist as both ds and ss DNA within the cell, it is the ssDNA form of the replicon that is packaged into the CP (Bennet and Agbandje-McKenna, 2020). Importantly, the host cell machinery expresses the viral genes using the dsDNA replicon as a template (Hanley-Bowdoin et al., 2013). Genes encoded in the BCTV virion-sense ORF are known to be involved in regulating the ratio of ss and ds DNA (Stanley and Latham, 1992; Hormuzdi and Bisaro, 1993). Since the *BCTVΔVS:YPET* construct lacks these genes, it is possible that the replicon preferentially accumulates in the dsDNA form in these deletion constructs, potentially explaining the enhanced YPET fluorescence.

A Southern blot was performed on DNA extracted from *N. benthamiana* leaves that were infiltrated with *Agrobacterium* carrying various BCTV-based viral vectors. Prior to Southern blotting, the DNA extract was treated with SalI restriction enzyme overnight to linearize the dsDNA. Both *BCTV:YPET* and *BCTVΔVS:YPET* contain a single SalI restriction site. The results of the Southern blot analysis demonstrated that although the *BCTV:YPET* and *BCTVΔVS:YPET* replicons accumulate at similar levels (Figure 3D), *BCTVΔVS:YPET* preferentially accumulates in a dsDNA form that is visible as a single 3-kb band (Figure 3E). On the other hand, *BCTV:YPET* produces a fainter 3-kb band and preferentially exists in the ssDNA form that is visible as a smearing pattern that migrates faster than the 3-kb band (Stanley and Latham, 1992). This difference in the ratio of ssDNA and dsDNA produced by the two vectors is another possible explanation for the difference in reporter gene expression and should be used to inform future vector design.

### Determining the size limit of the BCTV replicon

A major drawback of using viral vectors is that they can have restrictive cargo size limitations that prevent their use in certain applications. To determine the limits of our BCTV viral vector as well as to test our modular cloning approach, two BCTV-based constructs with large reporter genes were constructed: *BCTV:3xYPET* and *BCTV:RUBY*. *BCTV:3xYPET* contains a 2.2 kb triple *YPET* reporter composed of three individual *YPET* CDSs fused together (Zhou et al., 2011). The *BCTV:RUBY* construct contains a 4 kb *RUBY* reporter (He et al., 2020), a chimeric sequence that encodes three enzymes involved in the production of a reddish-purple pigment, betalain. These constructs were evaluated using *Agrobacterium*-mediated transient expression in *N. benthamiana*.

The *BCTV:3xYPET* and *BCTV:RUBY* constructs successfully mediated reporter expression (Figure 4A). When compared to a *35S:RUBY* construct delivered using a traditional T-DNA vector, the *BCTV:RUBY* vector resulted in stronger reporter expression. The *BCTVΔVS:RUBY* construct was also able to mediate strong expression of the *RUBY* reporter (Figure 4A). Given the size of these cargoes, it is possible that their inclusion in the viral vector has a negative impact on replicon accumulation. To test whether the inclusion of these large cargos negatively impacts replicon accumulation, qPCR was performed on DNA extracted from agroinfiltrated *N. benthamiana* leaves (Figure 4B). qPCR analysis demonstrated that there was no prominent difference in replicon accumulation between the *BCTV:YPET*, *BCTV:3xYPET, BCTV:RUBY,* and *BCTVΔVS:RUBY* vectors. This indicates that a cargo size of up to 4 kb does not have a significant impact on the ability of the replicon to accumulate within the cell. Although it remains unclear what the upper size limit may be, the fact that the BCTV replicon was able to handle the *RUBY* reporter indicates that our modular BCTV viral vectors can tolerate cargos of at least 4 kb in size, which is consistent with prior observations that cell-to-cell movement is the primary limiting factor for geminivirus genome size (Gilbertson et al., 2003).

**Figure 4.**
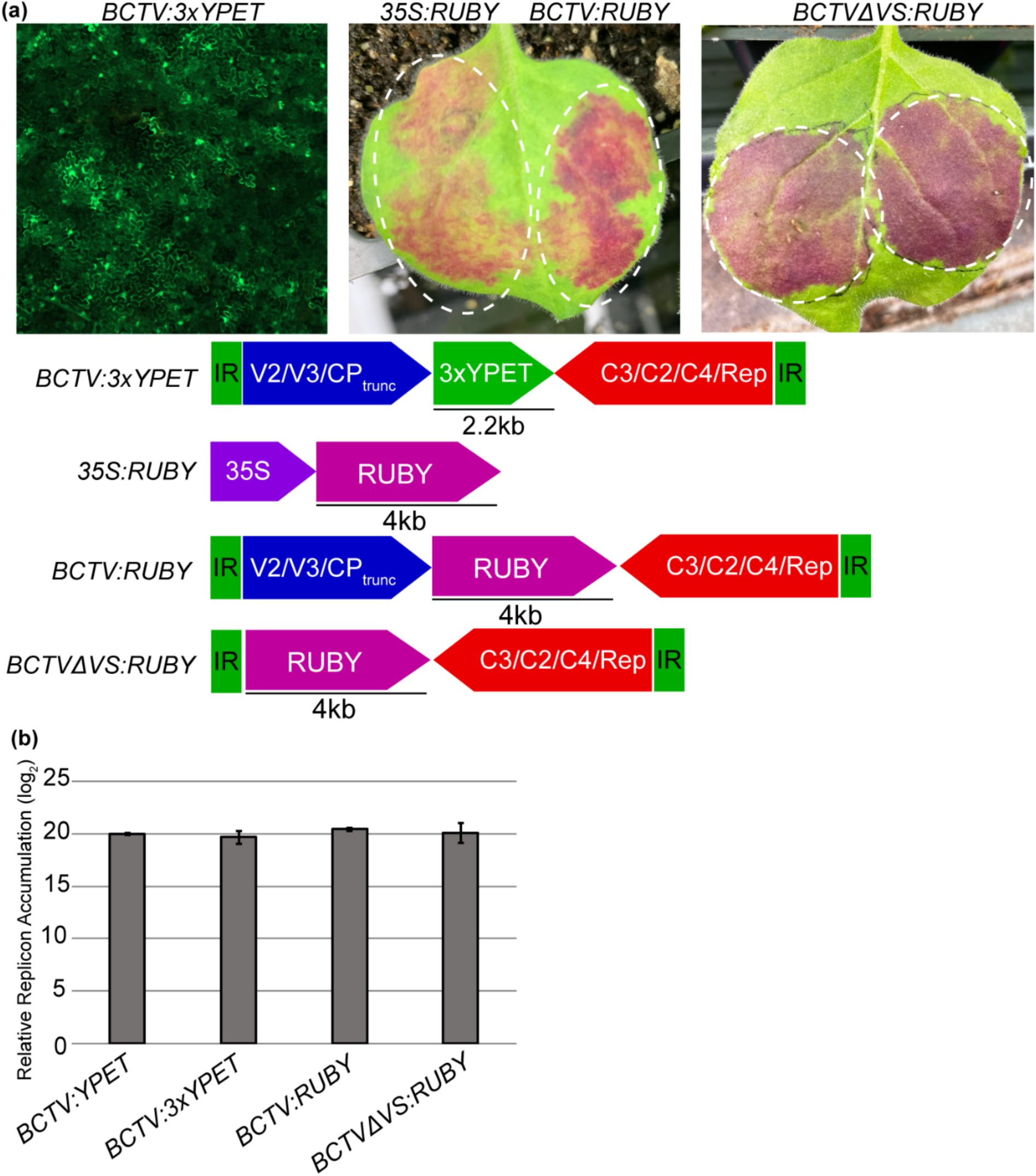
The BCTV viral vector can accommodate large cargos of at least 4kb. **A.** Fluorescence micrograph from a *N. benthamiana* leaf expressing a BCTV vector containing a triple *YPET* reporter (left). Leaves expressing the *RUBY* reporter under the control of the BCTV viral vector (left), the *35S* promoter (middle), or directly downstream of the left border-flanking IR (right). White dashed lines denote the infiltrated areas of the leaf. **B.** BCTV vectors with various cargoes accumulate to similar levels. qPCR was performed to determine whether the size of the cargo in the viral vector impacts BCTV accumulation. DNA was extracted from leaves at 5 dpi.

### Use of the BCTV viral vector in gene editing

Viral vectors, in addition to serving as useful tools for mediating high levels of protein expression, can also be leveraged as delivery vehicles for gene editing machinery. We sought to determine whether our BCTV-based vector could be used to deliver a repair template for GT, as well as whether the inclusion of the BCTV virion-sense genes has an impact on GT efficiency since these genes can affect the proportion of ss and ds viral DNA that accumulates in the plant cell (Hormuzdi and Bisaro, 1993).

To monitor HDR-based gene editing, we used a reporter line generated previously in *N. tabacum* (Wright et al., 2005). This reporter harbors a ’broken’ *GUS-NPTII* fusion that lacks part of the sequence necessary for enzymatic activity (Figure 5A). The reporter also contains a ZFN recognition site that can be cleaved to mediate gene editing. If successful GT occurs, the broken reporter is repaired, the inactive GUS enzyme is made active, and GT events can be detected by GUS staining.

**Figure 5.**
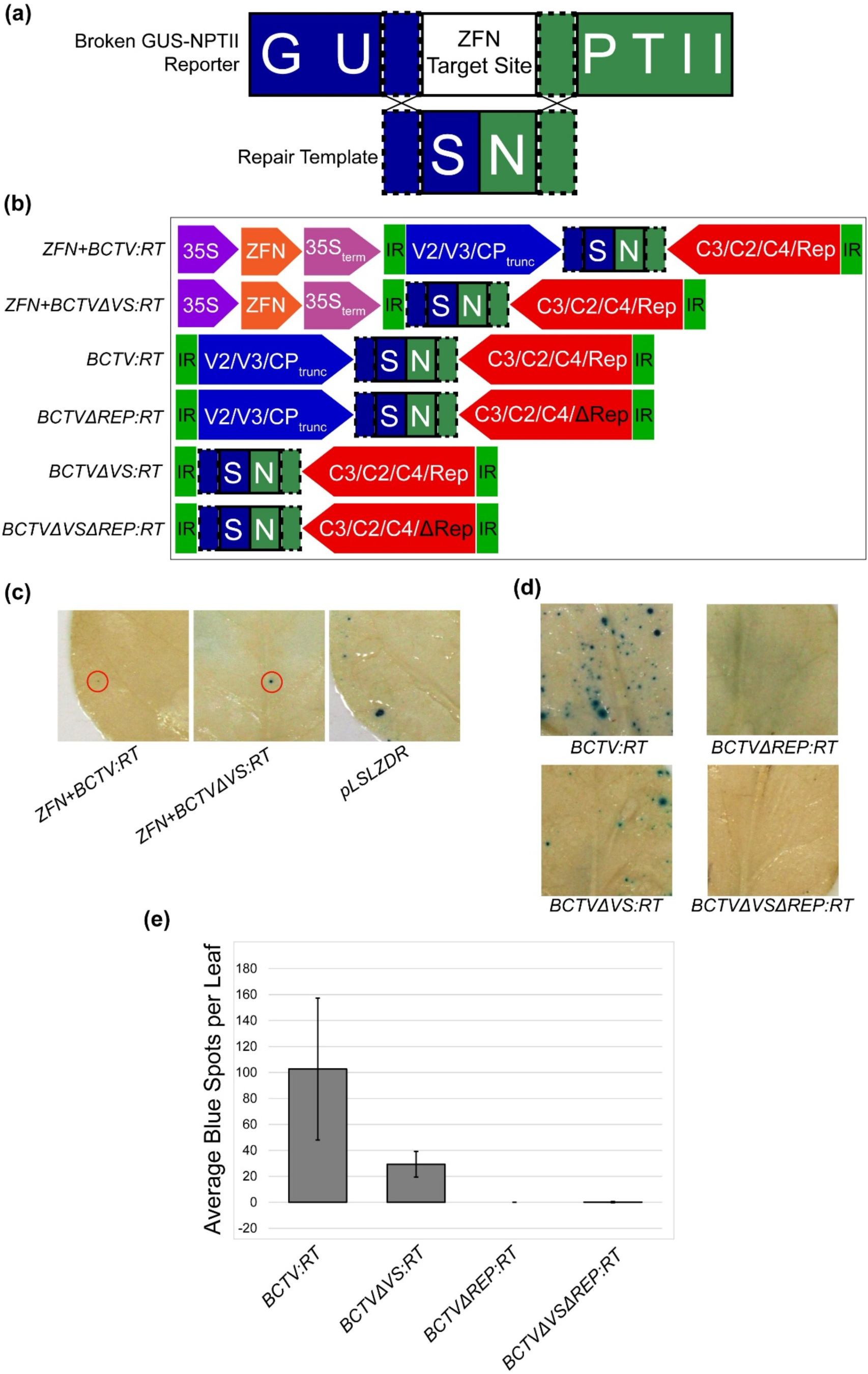
Optimization of the BCTV viral vector design for gene targeting (GT). **A.** Diagram of the broken *GUS* reporter construct used to detect successful GT events. The *GUS* reporter lacks sequence that is required for GUS activity, but it can be repaired if the repair template (RT) is used in the repair of the double-strand break induced by the ZFN. **B.** Diagram of the various BCTV constructs used for GT experiments. **C.** BCTV-delivered RT can successfully mediate GT in *N. tabacum*, but the BCTV-based constructs were less effective than the BeYDV-based pLSLZDR vector. Leaves were infiltrated with a single construct containing both a ZFN and a BCTV-delivered RT. Images of representative leaves subjected to GUS-staining are shown following chlorophyll removal, with a red circle indicating GUS-stained tissue. **D.** GT mediated by the BCTV viral vector requires *Rep* and is enhanced when the virion-sense genes are included. Leaves were co-infiltrated with a construct containing a *35S*-driven ZFN and the corresponding BCTV construct. Images of representative GUS-stained leaves are shown following chlorophyll removal. **E.** Quantification of successful GT events resulting from various BCTV constructs. Loss of *Rep* greatly reduced GT events. More successful GT events occurred when the BCTV virion-sense genes were included in the construct.

Two BCTV-based vectors containing a suitable RT were generated, *ZFN+BCTV:RT* and *ZFN+BCTVΔVS:RT* (Figure 5B). These vectors were assembled using GB cloning and contain a *35S*-driven ZFN that lies outside of the BCTV replicon sequences flanked by the *IR* motifs.

Thus, the ZFN is expressed under the control of the *35S* promoter contained within the T-DNA, while the RT accumulates within the BCTV replicon. The RT and ZFN used in these experiments were derived from the pLSLZDR Bean yellow dwarf virus (BeYDV) vector generated previously (Baltes et al., 2014). To test whether these new BCTV-based vectors can mediate successful GT, they were delivered into the broken *GUS* reporter *N. tabacum* line using *Agrobacterium*-mediated transient expression. After GUS staining, blue spots were observed, indicating that HDR took place. However, the efficiency of GT using the new BCTV-based vectors was typically lower than that observed when using the BeYDV-based pLSLZDR vector (Figure 5C). This suggests that BCTV may not function as well as BeYDV in *N. tabacum*.

We were interested in whether the BCTV-based vectors could function in other plant species in addition to *N. tabacum*. The ’broken’ *GUS:NPTII* reporter from *N. tabacum* was subcloned into pDGB3a1 and transformed into *N. benthamiana.* We selected two independent lines that performed well in GT experiments using the *ZFN+BCTV:RT* construct (Figure S3).

Interestingly, GT efficiency in these lines was much higher than that observed in the previously generated *N. tabacum* lines. This is likely attributable to *N. benthamiana* typically having a higher efficiency of *Agrobacterium*-mediated transient expression than *N. tabacum*. Using these *N. benthamiana* lines, we determined that BCTV-based viral vectors can deliver RT at high enough levels for efficient HDR-based gene editing (Figure 5D, E). We observed that the *ZFN+BCTV:RT* construct consistently mediated greater GT efficiency than the *ZFN+BCTVΔVS:RT* construct lacking the virion-sense genes. This suggests that although loss of the BCTV virion sense genes greatly enhances protein expression, it has the opposite effect on GT efficiency.

To test if successful BCTV-mediated gene targeting in *N. benthamiana* depends on the accumulation of the viral replicon, we generated two additional constructs that lack a functional *Rep* gene due to two in-frame stop codons in the 5’ end of the gene, *BCTVΔREP:RT* and *BCTVΔVSΔREP:RT*. As expected, loss of *Rep* prevented successful gene targeting events (Figure 5D, E), demonstrating that successful GT is dependent upon the copy number of the replicon-provided repair template.

## Discussion

Viral vectors have the potential to improve various fields of plant biotechnology, including gene editing and molecular farming. Successful gene targeting requires that a repair template and gene editing machinery be delivered to the cell, both of which can be mediated by viral vectors. Another promising use for viral vectors is in promoting high levels of protein expression. The use of plants to express important proteins is a potential way to produce pharmaceuticals and other valuable chemicals (Abrahamian et al., 2020). A greater understanding of how to design viral vectors can promote their use in plant biotechnology.

Previously generated viral vectors have generally been constructed using traditional cloning methods that can complicate the insertion of desired cargoes or alterations of the viral sequences. Here, we utilized a GB cloning approach to simplify the construct generation process. GB cloning has previously been used to create BeYDV-based viral vectors (Dusek et al., 2020). This approach enabled the authors to test different cargoes, but no alternative viral components were evaluated in that study. Since previous BCTV viral vector designs have included heterologous promoter sequences upstream of the cargo (Kim et al., 2007; Eini et al., 2022), we were interested in examining whether cargo expression is enhanced by these modifications. Interestingly, when we tested the inclusion of the widely used *35S* promoter upstream of the cargo, we found that it reduced cargo expression, despite having no effect on replicon accumulation (Figure 3C). Given that the *IR* motif contains elements that enable strong expression of the virion-sense genes, it is likely that the native elements mediate stronger expression than non-native elements like the *35S* promoter. In addition to improving cargo expression, the use of the native promoter elements also results in a smaller overall replicon size. Future vector designs, including those based on BCTV and other geminiviruses, should benefit from this observation.

BCTV machinery is involved in many different processes, including the replication of the viral replicon, manipulation of the host, and controlling the ratio of ds and ssDNA. We found that changes to the architecture of the viral vector, including the removal of the BCTV virion-sense genes, impact cargo expression. Removal of the BCTV virion-sense ORFs, including *V2* and *V3*, enhanced reporter expression while having no impact on replicon accumulation (Figures 3B, 3D). This may be due to the cargo being placed closer to the *IR* motif, which controls the expression of the virion-sense genes. We also observed that the removal of the BCTV virion-sense genes impacts the ratio of ds and ssDNA replicon that accumulates (Figure 3E). Thus, another possible explanation for the enhanced cargo expression in the BCTV clones lacking the virion-sense genes is that they provide more dsDNA to be transcribed by the host machinery.

Although we observed greater protein expression in BCTV constructs lacking the virion-sense genes, this architecture performed worse in GT experiments. A possible explanation for this is that the ssDNA replicon generated by constructs containing the virion-sense genes may serve as a more efficient template for GT than a dsDNA replicon. There is evidence to suggest that ssDNA templates may be more efficient for GT than dsDNA templates. In Chlamydomonas, Cas12-mediated GT was boosted 500-fold using ss oligodeoxynucleotides (ssODNs) (Ferenczi et al., 2017). Efficient GT using ssDNA templates in conjunction with a ZFN has also been demonstrated in human cell lines (Chen et al., 2011).

Another important consideration when using vectors derived from viruses is their cargo size limits. In general, the genome size, and therefore, the DNA cargo that viruses can efficiently pack into infectious particles, is typically limited by the requirements of plasmodesmata and the MP (Gilbertson et al., 2003). On the other hand, although the encapsidation restrictions would be irrelevant in virus-derived vectors lacking coat protein genes and unable to form infectious particles, other aspects of the virus biology may still impose certain cargo size limitations. For instance, large cargo sizes may affect the replication efficiency or the structural fidelity of the formed replicons. Here, we have shown that relatively large cargos of ∼4 kb do not significantly affect replicon accumulation or cargo protein expression, suggesting that most of the replicons formed contain the intact cargo sequence. We have also demonstrated that the deletion of the virion-sense genes does not have detrimental effects on cargo expression or replicon accumulation, suggesting that cargos even bigger than 4 kb could be accommodated in this BCTV-derived vector, making this new vector suitable for a large number of applications that require several kilobases of cargo DNA. It is also important to note that the stability, replication, and structural integrity of the replicons would likely depend on the specific sequences cloned in this vector. It would be interesting in future studies to test the effects of a broad range of DNA sizes and abundance of repetitive DNA on the replicon abundance and sequence stability.

Overall, these observations demonstrate the importance of designing a viral vector with its intended purpose in mind, whether that be to increase protein expression or deliver repair templates for gene targeting. The use of GB cloning enabled us to test numerous replicon architectures efficiently and optimize the BCTV viral vector for use in various applications. A similar approach could prove beneficial for designing and building more effective viral vectors for other plant viruses, host plants, and biotech applications.

## ACKNOWLEDGEMENTS

We would like to thank NC State University’s Plant Transformation Facility for their help with *N. benthamiana* transformation and the Genomic Sciences Laboratory for the sequencing services provided. We are also grateful to Dr. Diego Orzaez for sharing GB clones from his collection of standardized GB parts.

---

**Supporting Figure 1.**
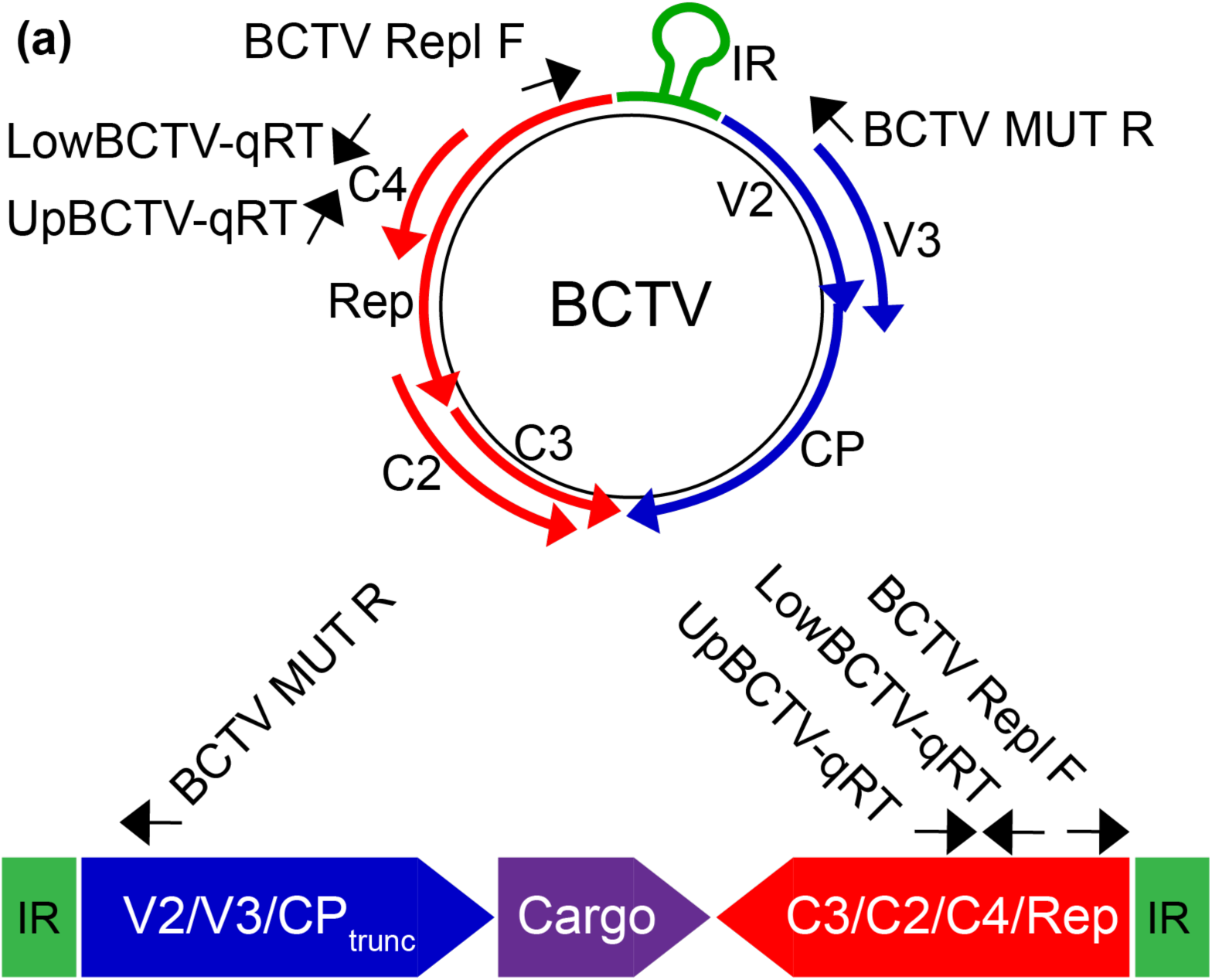
Diagram depicting the location of primers used in the study to detect the circularized BCTV replicon (BCTV Repl F and BCTV MUT R) and perform qPCR (UpBCTV-qRT and LowBCTV-qRT). BCTV Repl F and BCTV MUT R can only amplify the circular replicon, while UpBCTV-qRT and LowBCTV-qRT can amplify both the circular replicon and non-circularized viral vector.

**Supporting Figure 2.**
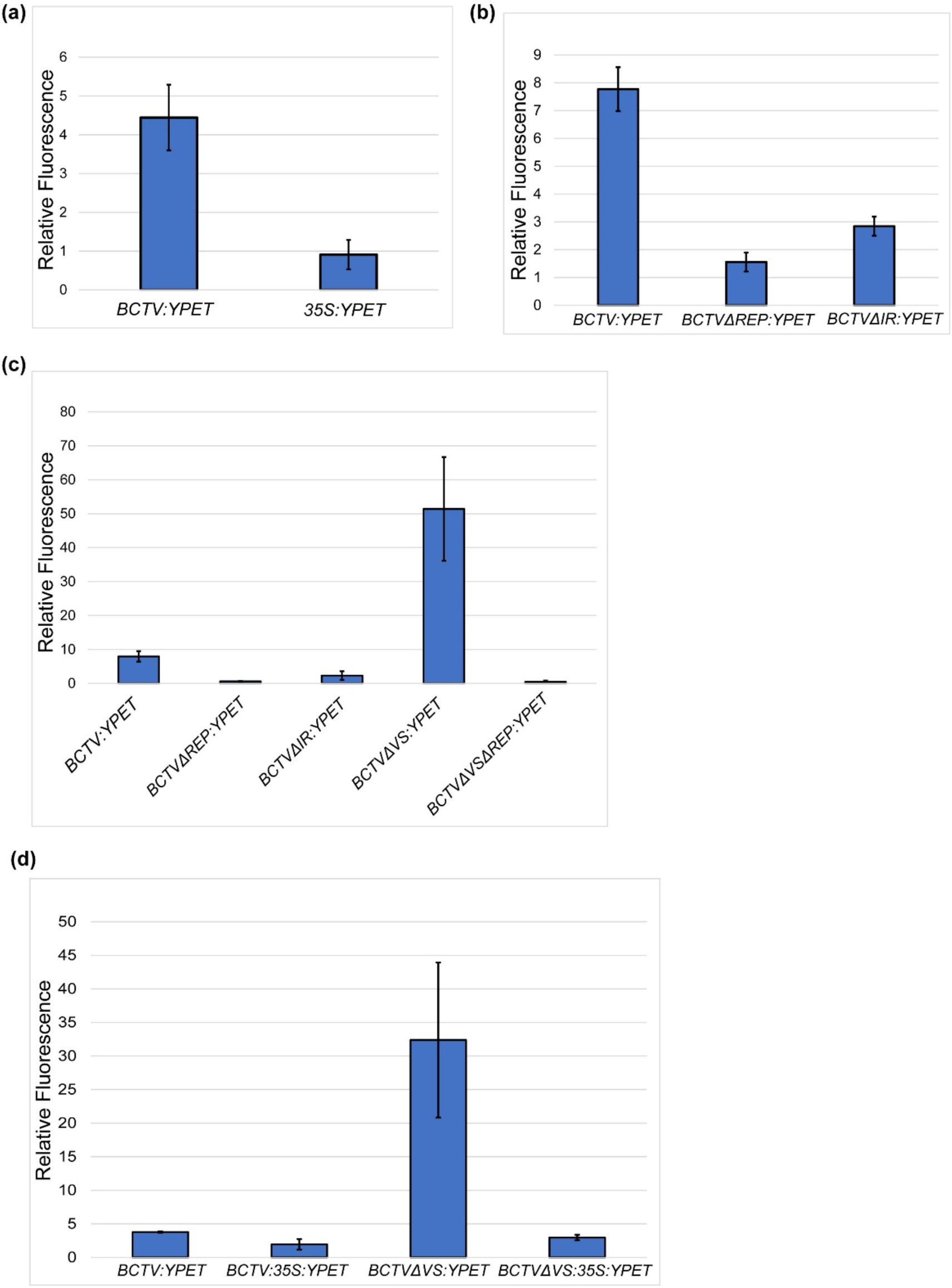
Quantification of fluorescence mediated by BCTV vectors harboring *YPET*. Image analysis was performed to compare fluorescence generated by various BCTV viral vectors. Averages were calculated based on three equal-size areas per each infiltration. Analysis was performed using ImageJ.

**Supporting Figure 3.**
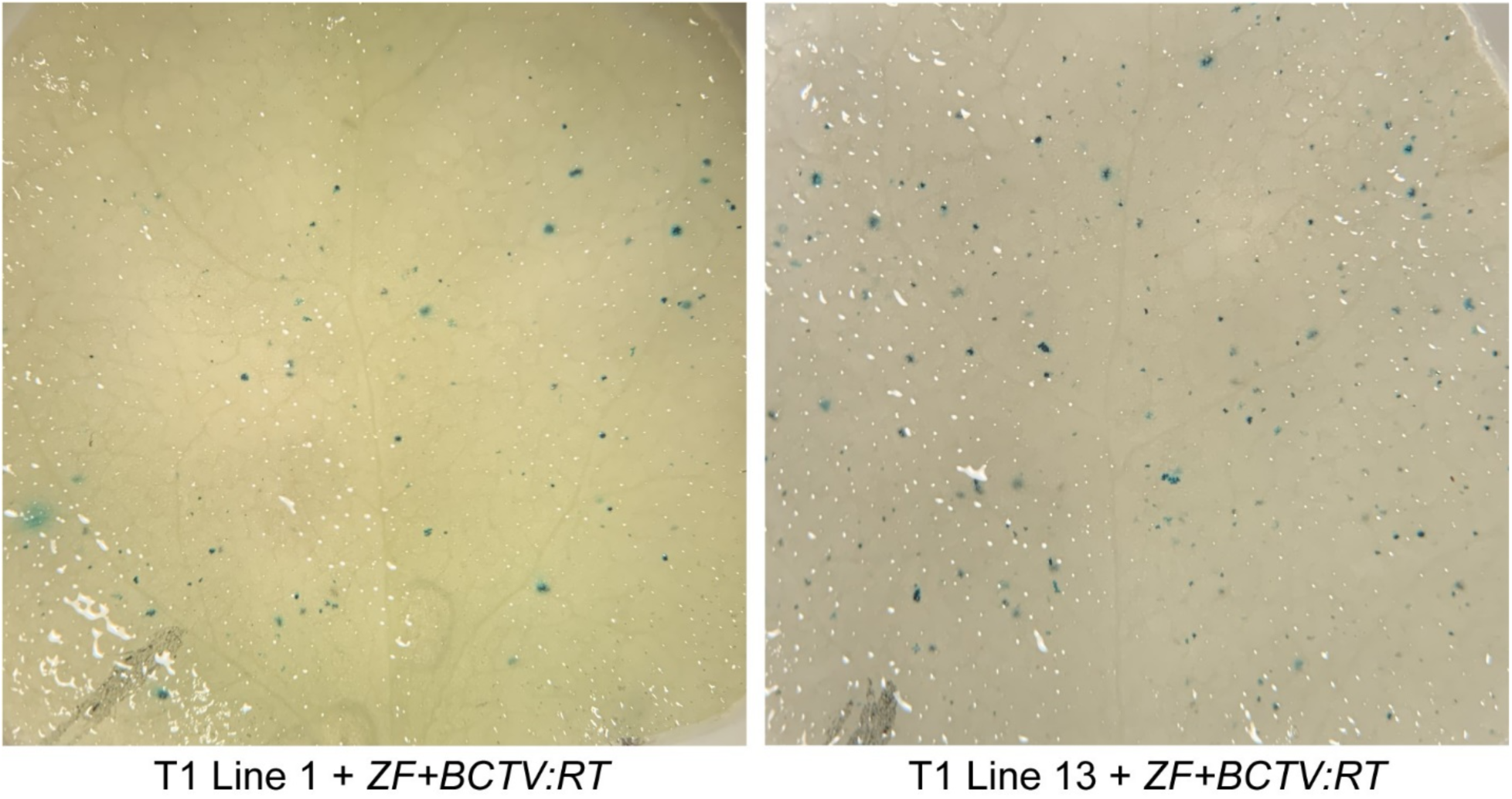
Two functional ‘broken’ *GUS* reporter lines were generated in *N. benthamiana*. Above, T1 plants from lines 1 and 13 were agroinfiltrated with strains carrying BCTV viral vectors with a zinc finger nuclease that targets the broken reporter and a repair template carrying the missing *GUS* sequence. Both lines consistently demonstrated successful repair of the *GUS* reporter and were used in subsequent experiments.

